# Prevention of tau accumulation through inhibition of hnRNP R-dependent axonal *Mapt* mRNA localization

**DOI:** 10.1101/2023.07.19.549639

**Authors:** Abdolhossein Zare, Saeede Salehi, Michael Briese, Michael Sendtner

**Author notes:** These authors contributed equally: Abdolhossein Zare, Saeede Salehi. These authors jointly supervised this work: Michael Briese, Michael Sendtner.

## Abstract

Deposition of neurofibrillary tangles composed of hyperphosphorylated tau in the brain is a pathological hallmark and closely correlates with onset and course of Alzheimeŕs disease. While tau reduction is being pursued as therapeutic strategy, prolonged lowering of total tau might lead to adverse effects, necessitating the development of more targeted approaches. We report that the RNA-binding protein hnRNP R facilitates the axonal localization of the *Mapt* mRNA encoding tau. Depletion of hnRNP R reduces tau in axons but not neuronal cell bodies. Brains of Alzheimer’s disease mice deficient for hnRNP R contain less tau tangles and amyloid-β plaques. Neurons treated with blocking antisense oligonucleotides to mask hnRNP R binding sites of *Mapt* mRNA show reduced axonal tau levels, similar to hnRNP R-deficient neurons. Lowering of tau levels selectively in axons, a major subcellular site of tangle formation and spreading, thus represents a novel therapeutic perspective for treatment of Alzheimer’s disease.

## Main

Alzheimer’s disease (AD) is a progressive neurodegenerative disorder primarily leading to memory loss but also affecting other cognitive functions over time. Pathologically, AD is characterized by the extracellular deposition of amyloid-β (Aβ) protein as senile plaques and the intracellular accumulation of neurofibrillary tangles containing the microtubule-associated protein tau. While Aβ plaques are present widespread in brains of AD patients, the temporal and spatial formation of tangles correlates more closely with the cognitive impairments and the progression of the disease^1–4^. Conversely, low tangle burden is associated with healthy brain aging^5^. Tau depletion prevents Aβ-induced neurotoxicity^6, 7^ and alleviates memory defects of AD mouse models^8, 9^, indicating a causal role of tau accumulation for AD pathogenesis. Thus, preventing tau pathology has become a main research avenue for AD therapy development^10^.

Current tau-targeting therapeutic strategies use antibodies, vaccines or antisense oligonucleotides (ASOs) to reduce total tau levels^10^. ASOs targeting the tau-encoding *MAPT* mRNA for RNase H-mediated degradation have proven effective in lowering tau protein levels in cultured neurons and mouse brain following intra-cerebroventricular administration^11, 12^. ASO-mediated tau reduction in a tauopathy mouse model expressing mutant P301S human tau prolonged survival and alleviated behavioural deficits^11^. However, globally reducing tau in the brain for prolonged times, particularly during aging, might have adverse consequences for neuronal functions. For example, while tau knockout mice appear phenotypically normal at middle age due to compensatory upregulation of other microtubule-associated proteins, they show synaptic defects and memory impairments at old age^13^. Additionally, acute tau knockdown in the hippocampus of 7-month-old mice causes motor and memory impairments accompanied by synaptic abnormalities^14^. Thus, more targeted approaches are needed to remove tau from subcellular sites at which tau pathology originates.

In AD, oligomeric and hyperphosphorylated tau accumulates in axons and at pre- and postsynaptic terminals^15, 16^. In the hippocampus of AD patients, axonal tau pathology precedes tau aggregation in somatodendritic regions, suggesting that formation of axonal tau deposits is an early event in the pathological AD cascade^17^. Beyond that, tau is secreted upon neuronal activation through interaction with synaptic vesicle proteins^18^, facilitating spreading of tau pathology between brain regions in AD^19–21^. Thus, targeted therapeutic approaches that selectively reduce axonal tau might prevent tangle formation and spreading while having less adverse effects compared to treatments that deplete tau globally.

In neurons, *MAPT* mRNA is transported into axons and locally translated into tau protein^22–24^. Thus, blocking axonal *MAPT* transport might reduce axonal tau protein production. Here, we show that the RNA-binding protein hnRNP R associates with the 3’ UTR of *Mapt*, and that loss of hnRNP R selectively reduces *Mapt* and tau levels in axons but not neuronal cell bodies. AD mice deficient in hnRNP R exhibit reduced plaque and tangle burden in the brain. We designed ASOs for blocking the association between hnRNP R and *Mapt* and observed a similar reduction in axonal *Mapt* and tau in neurons treated with these ASOs. Thus, ASO-mediated depletion of axonal tau might be a novel therapeutic strategy for treatment of AD and other tauopathies.

## Results

### Depletion of hnRNP R reduces axonal tau levels

We previously observed that the neuronal RNA-binding protein hnRNP R regulates the axonal localization of mRNAs in primary mouse embryonic motoneurons^25^. In line with this function, individual nucleotide resolution crosslinking and immunoprecipitation (iCLIP) revealed binding of cytosolic hnRNP R to 3’ UTRs of mRNAs^25^. To assess whether *Mapt* mRNA is targeted by hnRNP R, we visually inspected the *Mapt* locus and identified an enrichment of hnRNP R iCLIP hits in the 3’ UTR (Fig. 1a). RNA immunoprecipitation using an 4 antibody against hnRNP R confirmed its association with *Mapt* (Fig. 1b). Next, we tested whether depletion of hnRNP R alters the axonal localization of *Mapt*. For this purpose, we generated an hnRNP R knockout mouse model in which exons 1-5 of the *Hnrnpr* gene were deleted by CRISPR/Cas9-mediated genome engineering. Fluorescent *in situ* hybridization (FISH) revealed reduced axonal *Mapt* levels in motoneurons cultured from *Hnrnpr*^-/-^ mice relative to *Hnrnpr*^+/+^ motoneurons by (Fig. 1c,d). *Mapt* levels in the somata of *Hnrnpr*^-/-^ motoneurons were unchanged (Fig. 1d). Axonal, but not somatodendritic, levels of *Mapt* mRNA were also reduced in *Hnrnpr*^-/-^ compared to *Hnrnpr*^+/+^ motoneurons cultured in microfluidic chambers for 6 days *in vitro* (DIV) (Fig. 1e,f). In agreement with the FISH data, immunostaining revealed lower tau protein levels in axons but not cell bodies of *Hnrnpr*^-/-^ compared to *Hnrnpr*^+/+^ motoneurons (Fig. 1g,h). The effect of hnRNP R depletion on tau distribution was more prominent in distal versus proximal axons. This indicates that hnRNP R-mediated translocation of the *Mapt* mRNA to axons determines axonal tau protein levels.

**Fig. 1:**
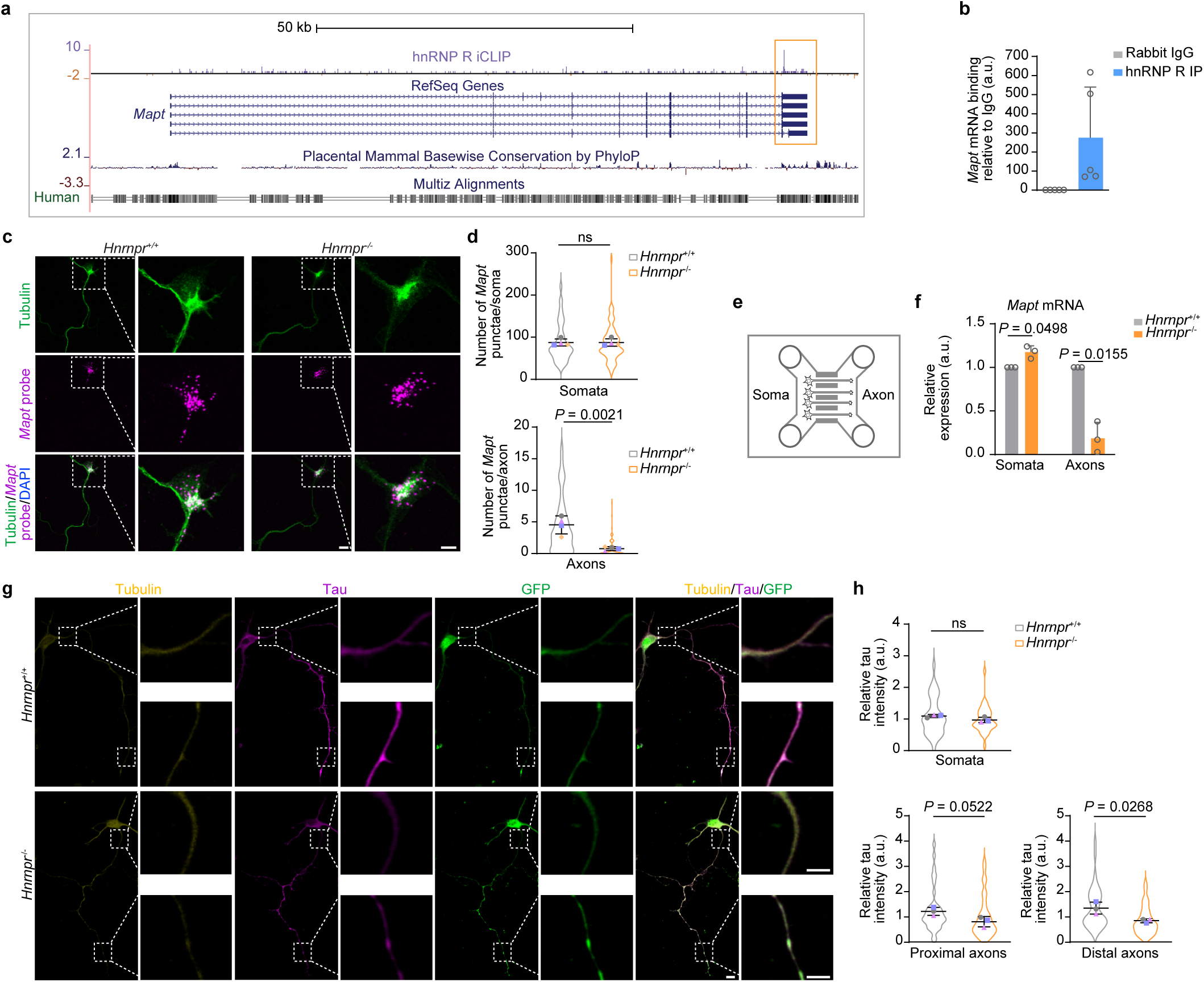
hnRNP R regulates axonal tau levels. **a**, UCSC genome browser view showing hnRNP R iCLIP hits along the *Mapt* pre-mRNA. **b**, Quantitative PCR (qPCR) of *Mapt* mRNA co-immunoprecipitated with an anti-hnRNP R antibody from DIV 7 mouse motoneurons. Data are mean ± standard deviation (s.d.) of *n*LJ=LJ5 independent experiments. **c**, Fluorescent *in situ* hybridization (FISH) of *Mapt* mRNA in DIV 5 motoneurons cultured from *Hnrnpr*^+/+^ and *Hnrnpr*^-/-^ mice. Scale bars: 10 µm and 5 µm (inset). **d**, SuperPlots of the number of *Mapt* FISH punctae in somata and axons of *Hnrnpr*^+/+^ and *Hnrnpr*^-/-^ motoneurons. Statistical analysis was performed using an unpaired two-tailed Student’s *t*-test. Data are mean ± s.d. of *n*LJ= 4 independent experiments; ns, not significant. **e**, Schematic of a microfluidic chamber for compartmentalized neuron cultures. **f**, qPCR analysis of *Mapt* mRNA from somatodendritic and axonal RNA of compartmentalized *Hnrnpr*^+/+^ and *Hnrnpr*^-/-^ mouse motoneurons at DIV 7. Statistical analysis was performed using a two-tailed one-sample *t*-test. Data are mean ± s.d. of *n*LJ=LJ3 independent experiments. **g**, Tau immunostaining of DIV 5 motoneurons cultured from *Hnrnpr*^+/+^ and *Hnrnpr*^-/-^ mice, with proximal and distal regions of the axon marked. Motoneurons were transduced with an EGFP expression lentivirus for visualization of neuronal morphology and for normalization of tau levels. Scale bars: 10 µm and 5 µm (inset). **h**, SuperPlots of tau immunointensities in the somata and proximal and distal axonal regions of *Hnrnpr*^+/+^ and *Hnrnpr*^-/-^ motoneurons. Statistical analysis was performed using an unpaired two-tailed Student’s *t*-test. Data are mean ± s.d. of *n*LJ=LJ3 independent experiments.

### Reduced AD pathology upon loss of hnRNP R

Our observation that hnRNP R depletion lowers axonal tau levels points towards the possibility that loss of hnRNP R ameliorates pathological alterations occurring in AD. To test this, we crossed *Hnrnpr^-/-^* with 5xFAD mice overexpressing mutant human amyloid-β precursor protein (APP) and mutant human presenilin-1 (PS1) harbouring familial AD mutations^26^. Brain sections of 6- and 9-months-old 5xFAD/*Hnrnpr*^+/+^ and 5xFAD/*Hnrnpr*^-/-^ mice were immunostained with the 6E10 antibody to visualize Aβ plaques. We observed a significant reduction in the number of Aβ plaques in the hippocampus and cortex of 5xFAD/*Hnrnpr*^-/-^ relative to 5xFAD/*Hnrnpr*^+/+^ mice at both ages (Fig. 2a,b). Deposition of Aβ plaques in brains of 5xFAD mice is accompanied by activation of microglia^27^. In agreement with the reduced plaque burden, we observed reduced microglial activation in the hippocampus and cortex of 5xFAD/*Hnrnpr*^-/-^ compared to 5xFAD/*Hnrnpr*^+/+^ mice following immunostaining for the microglial marker Iba1 (Fig. 2a,c). Additionally, immunostaining with 5 the AT8 antibody revealed significantly less tangles in these brain structures in 5xFAD/*Hnrnpr*^-/-^ mice (Fig. 2d,e). Depletion of hnRNP R thus alleviates AD pathology.

**Fig. 2:**
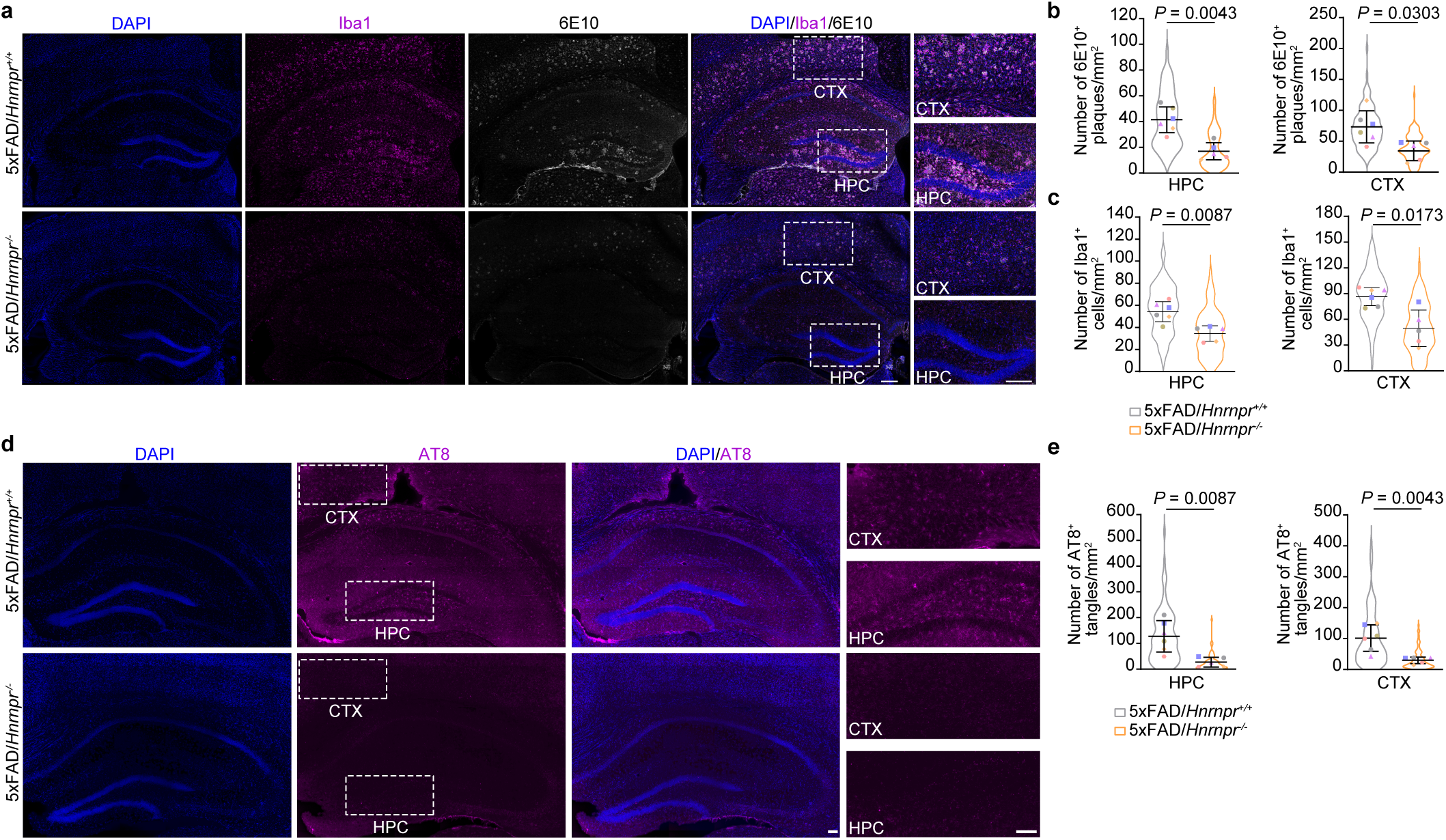
hnRNP R deficiency ameliorates tau and amyloid-β pathologies in 5xFAD mice. **a**, Immunohistochemical detection of microglia using an antibody against Iba1 and of Aβ plaques using the 6E10 antibody in coronal brain sections of 9-months-old 5xFAD/*Hnrnpr*^+/+^ and 5xFAD/*Hnrnpr*^-/-^ mice. CTX, cortex; HPC, hippocampus. Scale bars: 200 µm. **b**,**c**, SuperPlots of the number of Aβ plaques (**b**) and Iba1-positive microglia (**c**) per square millimetre in hippocampus and cortex of 6- and 9-months-old 5xFAD/*Hnrnpr*^+/+^ and 5xFAD/*Hnrnpr*^-/-^ mice. Statistical analysis was performed using an unpaired two-tailed Student’s *t*-test. Data are mean ± s.d. of *n*LJ=LJ6 5xFAD/*Hnrnpr*^+/+^ and *n* = 5 5xFAD/*Hnrnpr*^-/-^ mice. **d**, Immunohistochemical detection of hyperphosphorylated tau using the AT8 antibody in coronal brain sections of 9-months-old 5xFAD/*Hnrnpr*^+/+^ and 5xFAD/*Hnrnpr*^-/-^ mice. Scale bars: 200 µm. **e**, SuperPlots of the number of tau tangles per square millimetre in hippocampus and cortex of 6- and 9-months-old 5xFAD/*Hnrnpr*^+/+^ and 5xFAD/*Hnrnpr*^-/-^ mice. Statistical analysis was performed using an unpaired two-tailed Student’s *t*-test. Data are mean ± s.d. of *n*LJ=LJ6 5xFAD/*Hnrnpr*^+/+^ and *n* = 5 5xFAD/*Hnrnpr*^-/-^ mice.

### Antisense oligonucleotides for lowering axonal *Mapt* mRNA levels

Having shown that hnRNP R loss reduces axonal tau and lowers plaque and tangle deposition, we next sought to achieve axonal tau reduction more selectively through abolishing its localization into axons. For this purpose, we designed two 2’-*O*-methyl- and phosphorothioate-modified ASOs (MAPT-ASO1 and 2) complementary to *Mapt* 3’ UTR regions with hnRNP R iCLIP hits for blocking the association between hnRNP R and *Mapt* (Fig. 3a). We selected binding regions that were conserved between mouse and human. For analysis of uptake conditions, a Cy3-labeled sense oligonucleotide was used. Efficient uptake was observed by incubating motoneurons or hippocampal neurons with 10 µM oligonucleotide (Extended Data Fig. 1a,b), similar to a previous study^28^. Motoneurons treated with MAPT-ASO1 or 2 and cultured for 6 DIV showed reduced axonal *Mapt* mRNA levels compared to untreated motoneurons as detected by FISH (Extended Data Fig. 1c,d). A similar reduction in axonal *Mapt* was also detectable in cultured hippocampal neurons (Extended Data Fig. 1e,f). Importantly, *Mapt* levels in the cell bodies of MAPT-ASO-treated neurons were unchanged (Extended Data Fig. 1c-f).

**Fig. 3:**
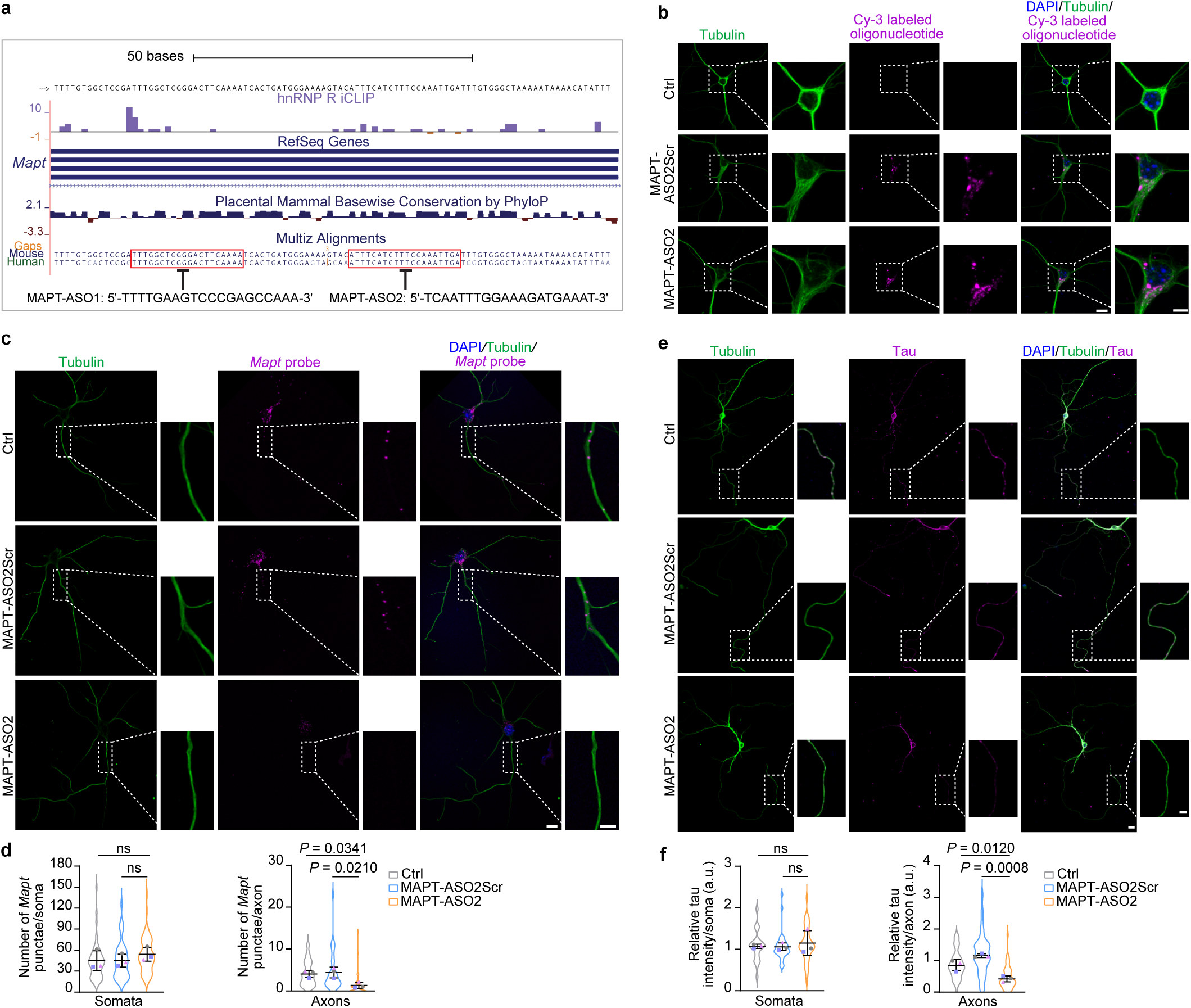
ASO-induced reduction of axonal tau. **a**, Design of two antisense oligonucleotides (MAPT-ASO1 and MAPT-ASO2) for blocking the interaction between hnRNP R and *Mapt* mRNA. **b**, Immunofluorescence imaging of DIV 25 untreated (Ctrl) mouse hippocampal neurons and hippocampal neurons treated with 10 µM of a Cy3-labeled scramble oligonucleotide or MAPT-ASO2. Scale bars: 10 µm and 5 µm (inset). Images are representative of at least three independent experiments. **c**, *Mapt* FISH of DIV 6 untreated (Ctrl) mouse hippocampal neurons and hippocampal neurons treated with scramble oligonucleotide or MAPT-ASO2. Scale bars: 10 µm and 5 µm (inset). **d**, SuperPlots of the number of *Mapt* FISH punctae in the somata and axons of hippocampal neurons. Statistical analysis was performed using a one-way ANOVA with Tukey’s multiple comparisons test. Data are mean ± s.d. of *n*LJ=LJ3 independent experiments. **e**, Tau immunostaining of DIV 25 untreated (Ctrl) mouse hippocampal neurons and hippocampal neurons treated with scramble oligonucleotide or MAPT-ASO2, with distal regions of the axon marked. Scale bars: 10 µm and 5 µm (inset). **f**, SuperPlots of tau immunointensities in the somata and distal axonal regions of hippocampal neurons. Statistical analysis was performed using a one-way ANOVA with Tukey’s multiple comparisons test. Data are mean ± s.d. of *n*LJ=LJ3 independent experiments.

Next, we investigated whether MAPT-ASO-mediated depletion of axonal *Mapt* can also downregulate tau protein in axons. Given the long half-life of tau^29^, we assessed tau levels by immunostaining in MAPT-ASO-treated as well as untreated motoneurons and hippocampal neurons cultured for 11 and 22-25 DIV, respectively. Following MAPT-ASO treatment, tau levels were reduced in axons but not cell bodies of motoneurons or hippocampal neurons (Extended Data Fig. 2a-d). The axonal tau reduction was more pronounced for MAPT-ASO2 compared to MAPT-ASO1 (Extended Data Fig. 2b,d). Together, these data indicate that tau protein levels can be selectively reduced in axons by ASOs blocking the hnRNP R binding sites in the *Mapt* 3’ UTR.

### MAPT-ASO2 reduces axonal tau

Since MAPT-ASO2 more efficiently reduced axonal tau relative to MAPT-ASO1, we used MAPT-ASO2 and compared it to a scrambled version of it (MAPT-ASO2Scr) as additional control. At 10 µM, Cy3-labeled MAPT-ASO2 and Scr were efficiently taken up by motoneurons (Extended Data Fig. 3a) and hippocampal neurons (Fig. 3b). Compared to Scr, MAPT-ASO2 significantly downregulated *Mapt* levels in axons but not somata of motoneurons (Extended Data Fig. 3b,c) and hippocampal neurons (Fig. 3c,d). Likewise, tau protein levels were reduced in axons of MAPT-ASO2-treated motoneurons (Extended Data Fig. 3d,e) and hippocampal neurons (Fig. 3e,f) compared Scr-treated neurons. In contrast, tau levels in the somata of MAPT-ASO2-treated neurons remained unchanged (Fig. 3e,f and Extended Data Fig. 3d,e). We then used a puromycin labelling-proximity ligation assay (Puro-PLA)^30^ to assess the axonal translation of tau. Motoneurons treated with MAPT-ASO2 revealed a reduced Puro-PLA signal for tau in axons compared to Scr-treated and untreated motoneurons (Extended Data Fig. 4a,b). Importantly, tau synthesis in cell bodies was unaffected by MAPT-ASO2 treatment (Extended Data Fig. 4b). Thus, MAPT-ASO2 treatment can reduce axonal levels of *Mapt*, resulting in less axonal tau due to lowered local translation. In agreement with reduced axon growth observed in tau-depleted neurons^31, 32^, axon lengths of motoneurons subjected to MAPT-ASO2 treatment were reduced compared to Scr treatment while survival was unaffected (Fig. 4a-c). Reduced axon growth was also detectable for hippocampal neurons exposed to MAPT-ASO2 (Fig. 4d,e). In hippocampal neurons, survival remained unchanged following MAPT-ASO2 treatment (Fig. 4f). MAPT-ASO2 treatment therefore phenocopies tau deficiency.

**Fig. 4:**
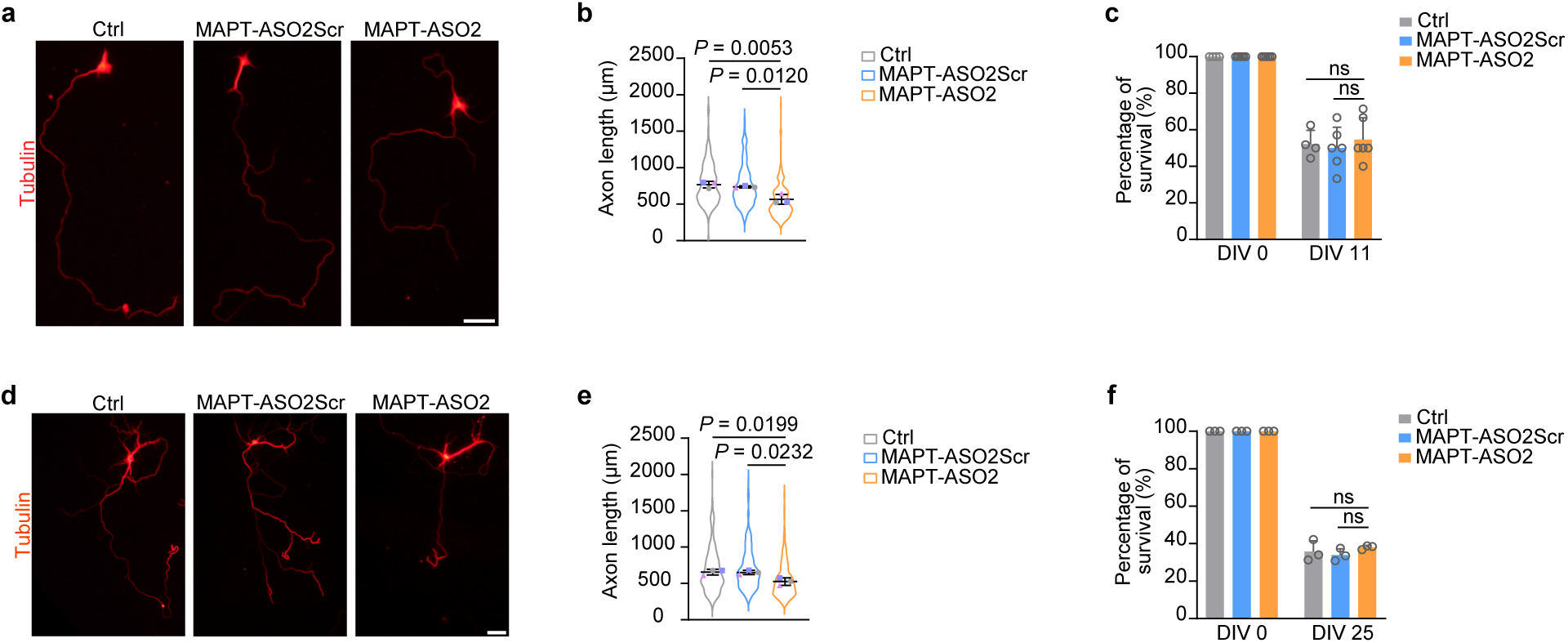
Axonal tau reduction phenocopies tau deficiency. **a**, Morphology of DIV 7 untreated (Ctrl) mouse motoneurons and motoneurons treated with scramble oligonucleotide or MAPT-ASO2. Scale bar: 50 µm. **b**, SuperPlots of axon lengths of motoneurons. Statistical analysis was performed using a one-way ANOVA with Tukey’s multiple comparisons test. Data are mean ± s.d. of *n*LJ=LJ3 independent experiments. **c**, Quantification of motoneuron survival on DIV 11. Statistical analysis was performed using a two-way ANOVA with Tukey’s multiple comparisons test. Data are mean ± s.d. of *n*lJ=LJ6 independent experiments. **d**, Morphology of DIV 25 untreated (Ctrl) mouse hippocampal neurons and hippocampal neurons treated with scramble oligonucleotide or MAPT-ASO2. Scale bar: 50 µm. **e**, SuperPlots of axon lengths of hippocampal neurons. Statistical analysis was performed using a one-way ANOVA with Tukey’s multiple comparisons test. Data are mean ± s.d. of *n*LJ=LJ3 independent experiments. **f**, Quantification of hippocampal neuron survival on DIV 25. Statistical analysis was performed using two-way ANOVA with Tukey’s multiple comparisons test. Data are mean ± s.d. of *n*lJ=LJ3 independent experiments.

Next, we designed additional MAPT-ASOs along the *Mapt* 3’ UTR in regions that contain hnRNP R iCLIP hits and that are conserved between mouse and human (Extended Data Fig. 5a,b). We screened these ASOs by FISH in hippocampal neurons for their potential to reduce axonal *Mapt* mRNA levels and identified several candidates that lowered *Mapt* levels to less than 50% relative to untreated hippocampal neurons (Extended Data Fig. 5b,c). Of these, MAPT-ASO5 overlapped the binding site of MAPT-ASO1 and MAPT-ASO19 and 20 were shortened versions of MAPT-ASO2 with a length of 18 and 16 nucleotides, respectively (Extended Data Fig. 5a,b). Treatment of hippocampal neurons with MAPT-ASO5 or 20 reduced axonal levels of *Mapt* mRNA, tau protein and axon growth similar to treatment with MAPT-ASO1 and 2 (Fig. 5a-d). Taken together, our findings indicate that the *Mapt* 3’ UTR can be targeted by blocking ASOs to reduce tau levels locally in axons.

**Fig. 5:**
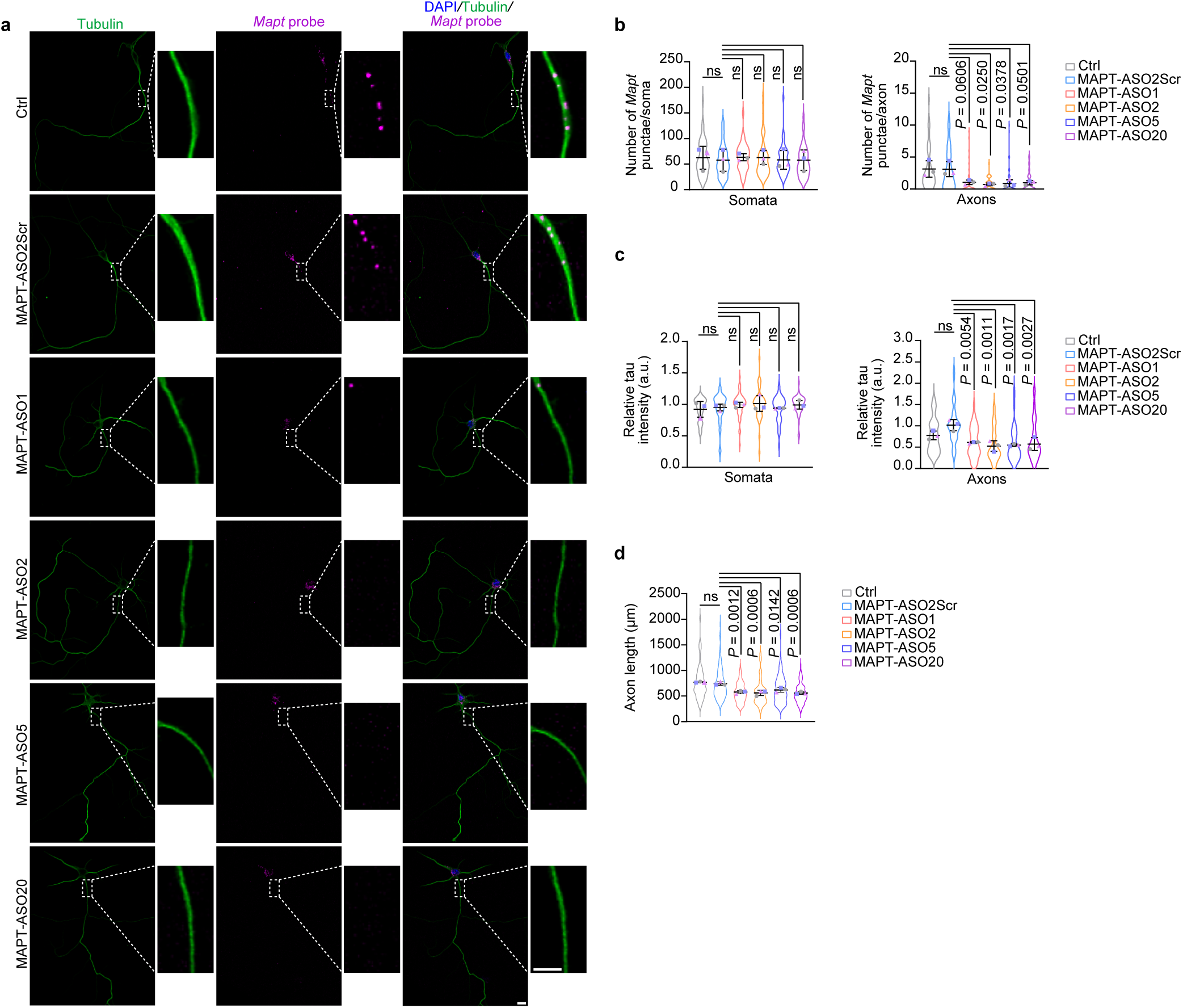
Identification of additional ASOs for axonal tau reduction. **a**, *Mapt* FISH of DIV 6 untreated (Ctrl) mouse hippocampal neurons and hippocampal neurons treated with scramble oligonucleotide or MAPT-ASO1, 2, 5 or 20. Scale bars: 10 µm and 5 µm (inset). **b**, SuperPlots of the number of *Mapt* FISH punctae in the somata and axons of hippocampal neurons. Statistical analysis was performed using a one-way ANOVA with Tukey’s multiple comparisons test. Data are mean ± s.d. of *n*LJ=LJ3 independent experiments. **c**, SuperPlots of tau immunointensities in the somata and distal axonal regions of DIV 25 hippocampal neurons. Statistical analysis was performed using a one-way ANOVA with Tukey’s multiple comparisons test. Data are mean ± s.d. of *n*LJ=LJ3 independent experiments. **d**, SuperPlots of axon lengths of DIV 25 hippocampal neurons. Statistical analysis was performed using a one-way ANOVA with Tukey’s multiple comparisons test. Data are mean ± s.d. of *n*LJ=LJ3 independent experiments.

### ASO-mediated reduction of axonal tau in iPSCs-derived human motoneurons

We then tested MAPT-ASO2 for its efficacy in reducing axonal tau in human neurons. For this purpose, we differentiated induced pluripotent stem cells (iPSCs) into cholinergic human motoneurons using small molecules^33^ (Fig. 6a). Motoneurons were treated with 10 µM MAPT-ASO2 or Scr, or left untreated. At this concentration, ASO was detectable in motoneurons cultured for 25 DIV (Fig. 6b). Compared to untreated motoneurons, or motoneurons exposed to MAPT-ASO2Scr, treatment with MAPT-ASO2 reduced axonal tau levels, both in proximal and distal regions (Fig. 6c,d). In contrast, tau levels in the somata of MAPT-ASO2-treated motoneurons were unchanged (Fig. 6c,d). Thus MAPT-ASO2 lowers axonal tau in human neurons.

**Fig. 6:**
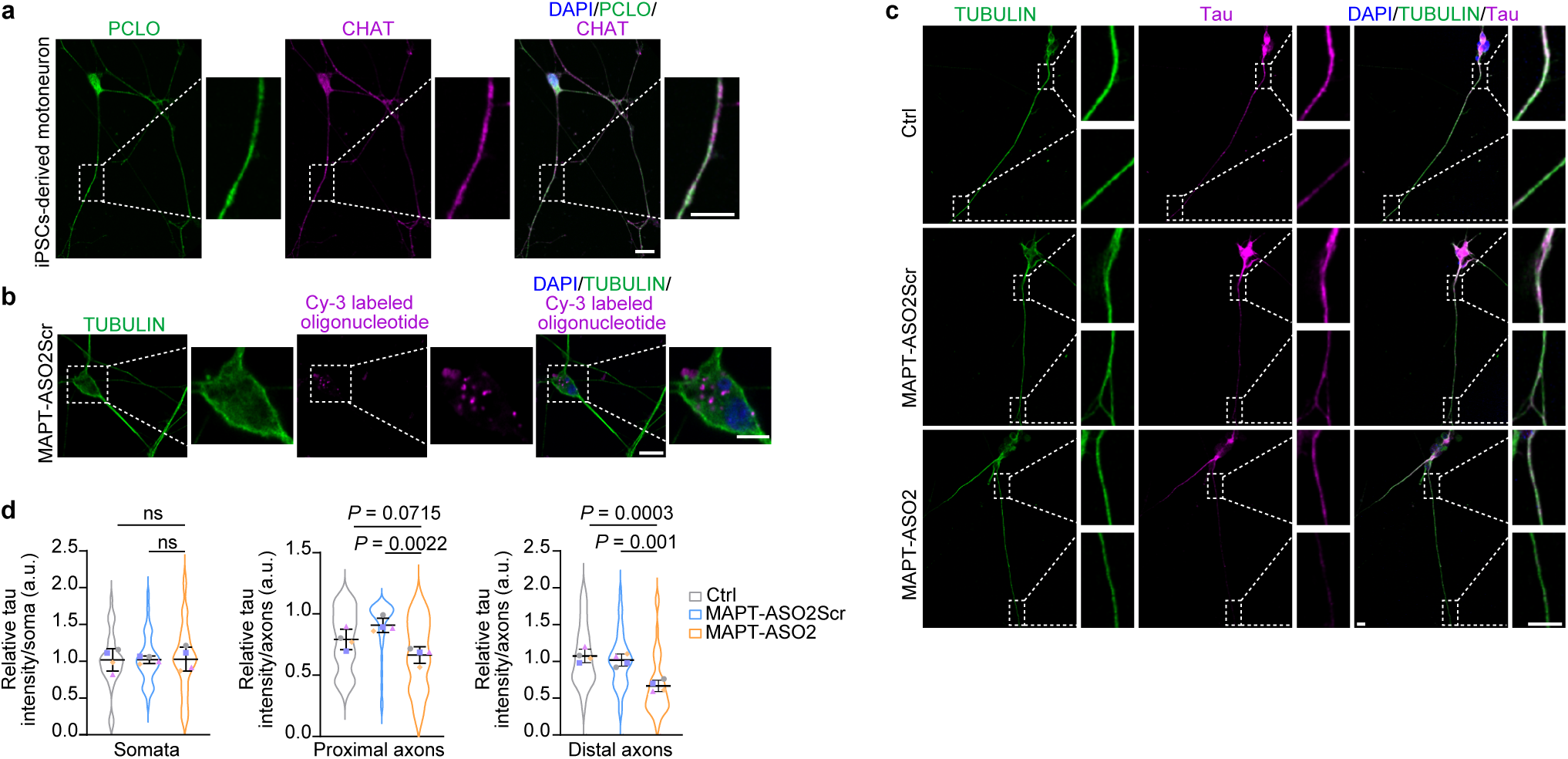
ASO-mediated reduction of axonal tau in iPSCs-derived human motoneurons. **a**, Immunostaining of DIV 25 iPSC-derived human motoneurons for PICCOLO (PCLO) and choline acetyltransferase (CHAT). Scale bars: 10 µm and 5 µm (inset). **b**, Immunofluorescence imaging of DIV 25 iPSC-derived human motoneurons treated with 10 µM of a Cy3-labeled scramble oligonucleotide or MAPT-ASO2. Scale bars: 10 µm and 5 µm (inset). Images in **a** and **b** are representative of at least three independent experiments. **c**, Tau immunostaining of DIV 25 untreated (Ctrl) iPSC-derived human motoneurons and iPSC-derived human motoneurons treated with scramble oligonucleotide or MAPT-ASO2, with proximal and distal regions of the axon marked. Scale bars: 10 µm and 5 µm (inset). **d**, SuperPlots of tau immunointensities in the somata and proximal and distal axonal regions of iPSC-derived human motoneurons. Statistical analysis was performed using a one-way ANOVA with Tukey’s multiple comparisons test. Data are mean ± s.d. of *n*LJ=LJ4 independent experiments.

## Discussion

Tau reduction has emerged as a promising therapeutic approach for treatment of AD and other tauopathies. Several strategies are currently being developed to lower tau levels. Immunotherapies utilize antibodies against tau to induce clearance of tangles. However, the existence of multiple tau splice isoforms and oligomeric tau species requires careful identification of the most promising tau form to be targeted^34^. Additionally, immunotherapies might be associated with increased risks for adverse effects such as inflammatory responses, cerebral edemas or hemorrhages. An alternative thus might be the depletion of tau through ASO-mediated degradation of the *MAPT* mRNA. While this approach can lower tau levels in the nervous system, leading to amelioration of symptoms in tauopathy mouse models, prolonged depletion of tau throughout neurons might be detrimental. Thus, the development of more targeted approaches to prevent tau elevation and tangle formation selectively in axons while sparing physiological functions of tau in neuronal cell bodies might prevent the occurrence of side effects associated with global tau depletion.

The strategy outlined here makes use of the hnRNP R-dependent axonal transport of *Mapt* mRNA and its local translation into tau protein. By blocking the interaction between *Mapt* and hnRNP R with ASOs, a reduction in axonal tau levels can be achieved without affecting tau levels in the somatodendritic compartment (Fig. 7). While we characterized MAPT-ASO2 in detail, other ASOs described here might be equally efficient for lowering axonal tau. Interestingly, the efficacy of MAPT-ASO19 and 20 was comparable to that of MAPT-ASO2. Since 19 and 20 are shorter versions of MAPT-ASO2, this finding points towards this particular *Mapt* 3’ UTR region as the most promising target sequence. Together, our findings represent an important step towards more selective tau-targeting strategies. If administered sufficiently early in the course of the disease, the MAPT-ASOs described here might be useful for limiting the initiation and spreading of tau pathology in AD in a more targeted manner compared to approaches aimed at global tau depletion. Either alone or in combination with recently developed therapies targeting Aβ plaques^35^, axonal tau-targeting MAPT-ASOs might thus hold the potential to slow down AD progression.

**Fig. 7:**
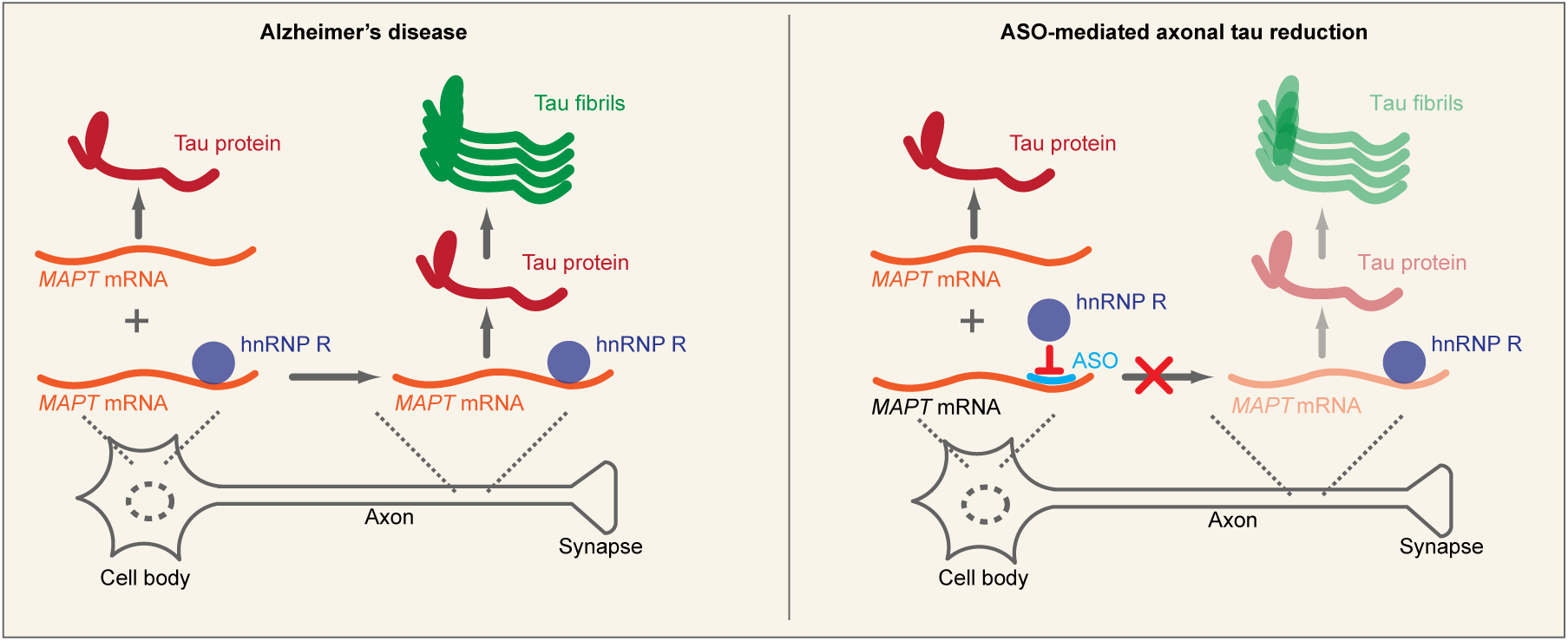
Schematic representation of ASO-mediated reduction in axonal tau levels.

## Methods

### Animals and ethical approval

The experimental procedures in this study involving mice were performed according to the regulations on animal protection of the German federal law and the Association for Assessment and Accreditation of Laboratory Animal Care, in agreement with and under the control of the local veterinary authority. Mice were housed in the animal facility of the Institute of Clinical Neurobiology at the University Hospital of Wuerzburg. Mice were maintained on a 12 h/12 h day/night cycle under controlled conditions at 20-22°C and 55-65% humidity with food and water in abundant supply. 5xFAD transgenic mice (034848-JAX) overexpressing human APP with the familial AD mutations K670N/M671L, I716V, and V717I and human PS1 with the familial AD mutations M146L and L286V were obtained from The Jackson Laboratory. *Hnrnpr*^-/-^ mice were generated by CRISPR/Cas9-mediated removal of a genomic region encompassing *Hnrnpr* exons1-5 using the gRNA target sequences 5’-ATATGCATAGTCTACTGCCCTGG-3’ and 5’-CCCAGACTACCTTGACGATGGGG-3’. A detailed description of *Hnrnpr*^-/-^ mice will be published elsewhere.

### Primary embryonic mouse motoneuron culture

**I**solation and enrichment of primary mouse motoneurons were performed as previously described^36^. Briefly, lumbar spinal cords were isolated from E13 mouse embryos, and motoneurons were enriched by p75NTR antibody (clone MLR2, Biosensis) panning. Motoneurons were plated on coverslips or in culture dishes coated with poly-DL-ornithine hydrobromide (PORN; P8638, Sigma-Aldrich) and laminin-111 (23017-015, Thermo Fisher Scientific). Motoneurons were maintained at 37°C, 5% CO_2_ in neurobasal medium (Gibco) supplemented with 2% B27 (Gibco), 2% heat-inactivated horse serum (Linaris), 1% GlutaMAX (Gibco) and 5 ng/mL of brain-derived neurotrophic factor (BDNF). Medium was replaced one day after plating and then every second day. For compartmentalization, motoneurons were grown in microfluidic chambers (Xona Microfluidics, SND 150) using a BDNF gradient as described before^37^. For qPCR analysis of compartmentalized motoneurons, total RNA was extracted using the PicoPure RNA Isolation Kit (KIT0204, Thermo Fisher Scientific) and reverse-transcribed using the First Strand cDNA Synthesis Kit (K1612, Thermo Fisher Scientific).

### Hippocampal neuron culture

Hippocampi dissected from P1 mice were immersed in dissection medium (Hanks’ Balanced Salt Solution (HBSS; Gibco) containing 1 mM sodium pyruvate (Gibco), 0.1% glucose and 10 mM HEPES pH7.3) containing 0.1% Trypsin (Worthington Biochemical Corporation) for 20 min at 37°C. Hippocampi were washed three times with dissection medium followed by addition of DNase (0.1%; Sigma-Aldrich) and incubation for 5 min at room temperature. After two washes with dissection medium, trypsin inhibitor (0.1%; Sigma-Aldrich) was added and hippocampi were triturated. Cells were centrifuged at 300 × g for 3 min, washed in dissection medium followed by centrifugation and resuspension neurobasal medium containing 2% B27, 1% penicillin/streptomycin, 1% GlutaMAX and 1% heat-inactivated horse serum. Cells were plated on poly-L-lysine-(P2636, Sigma-Aldrich) and laminin-111-coated glass coverslips for 1 h followed by incubation in neurobasal medium containing 2% B27, 1% penicillin/streptomycin and 1% GlutaMAX for 6–25 DIV. Half of the medium was replaced every four days.

### Differentiation of iPSC-derived motoneurons

For the generation of iPSC-derived motoneurons, we used a previously reported human iPSC-line 34D6 (control iPSC-line #1)^38^, which was kindly provided by Dr. Selvaraj and Prof. Chandran (University of Edinburgh). Motoneurons were differentiated according to a previously published protocol with some modifications^33^. Briefly, iPSCs were expanded in mTeSR Plus medium (05825, Stemcell Technologies) on Matrigel-coated (356234, Corning) dishes. Cells were passaged at 70% confluency using ReLeSR reagent (05872, Stemcell Technologies). After passaging, cells were cultured in the presence of ROCK Inhibitor (10 µM; 130-106-538, Miltenyi Biotec) for 24 h. For initiation of neuronal induction, iPSCs were maintained in mTeSR Plus medium supplemented with 10 µM SB431542 (13031, Cayman Chemical Company), 1 µM dorsomorphin homolog 1 (4126, R&D Systems), 3 µM CHIR99021 (13122, Cayman Chemical Company) and 0.5 µM purmorphamine (PMA; 10009634, Cayman Chemical Company). On day 2, medium was replaced by N2B27 medium containing Neurobasal medium (Gibco), Dulbecco’s modified Eagle’s medium F-12 (DMEM/F-12; Gibco), MACS NeuroBrew-21 (130-097-263, Miltenyi Biotech), N-2 Supplement (17502048, Gibco), 1% penicillin-streptomycin-glutamine supplemented with the same small molecules. On day 4, medium was replaced by expansion medium containing N2B27 medium supplemented with 3 µM CHIR99021, 0.5 µM PMA and 150 µM ascorbic acid (A4544, Sigma-Aldrich). Confluent cells were passaged and maintained in suspension on uncoated dishes. Embryoid bodies (EBs) were picked and triturated with a 1 mL pipette into smaller pieces on day 6. Subsequently, the cells were plated in expansion medium on Matrigel-coated dishes. The resulting neural precursor cells (NPCs) were passaged once every week with accutase (A1110501, Thermo Fisher Scientific) and expanded for at least 15 passages in expansion medium. Medium was replaced every other day. To induce motoneuron patterning, NPCs were kept for 9 days in N2B27 medium supplemented with 1 µM PMA. On day 2, 1 µM retinoic acid (72264, Stemcell Technologies) was added to the medium. The medium was replaced every other day. For differentiation, the medium was replaced with N2B27 medium supplemented with 10 ng/mL glia-derived neurotrophic factor (G-240, Alomone Labs), 10 ng/mL brain-derived neurotrophic factor (167450-02-STD, tebu-bio) and 500 µM dibutyryl-cAMP (Stemcell Technologies, 73886) and motoneurons were maintained for 25 DIV.

### Antisense oligonucleotides (ASOs)

ASOs were synthesized with full phosphorothioate backbone and 2’-O-Methyl groups at all nucleotides (Metabion). For ASO treatment, cells were incubated with 10 µM ASO or Scramble control oligonucleotide for 15 min at 37°C prior to plating. Mouse hippocampal and motoneurons were treated with ASO on DIV 0 and human neurons were treated prior to differentiation.

### RNA immunoprecipitation

For RNA immunoprecipitation, primary mouse motoneurons were grown on laminin-111-coated 6 cm dishes for 7 DIV. Cells were washed once with Dulbecco’s Phosphate Buffered Saline (DPBS, without MgCl_2_, CaCl_2_; D8537, Sigma-Aldrich) and lysed in lysis buffer (20 mM Tris-HCl pH7.4, 150 mM KCl, 2LJmM MgCl_2_, 0.1% NP-40). Lysates were incubated on ice for 20 min and centrifuged at 20,000 × g for 10 min at 4°C. Magnetic Dynabeads protein A (Thermo Fisher Scientific) were bound to 1 μg hnRNP R antibody (ab30930, Abcam) or 1 μg rabbit IgG control in 300 µl wash buffer (10 mM HEPES pH7.0, 100 mM KCl, 5LJmM MgCl_2_, 0.5% NP-40) for 1 h at room temperature by rotation. Antibody-bound beads were washed twice with wash buffer, and 300 μl lysate was added to the beads and incubated for 2 h at 4°C by rotation. Beads were washed twice with 500 µl wash buffer. Total RNA was extracted from the input sample and beads by adding 300 μl buffer A1 (NucleoSpin RNA kit, Macherey-Nagel) and 300 μl absolute ethanol followed by RNA extraction according to the manufacturer’s instructions. RNA was reverse-transcribed with random hexamers using the First Strand cDNA Synthesis Kit (Thermo Fisher Scientific). Reverse transcription reactions were diluted 1:5 in water and transcript levels were measured by qPCR. Relative RNA binding was calculated using the ΔΔCt method with normalization to input levels.

### Fluorescence *in situ* hybridization (FISH)

FISH was performed with the ViewRNA ISH Cell Assay kit (QVC0001, Invitrogen). Motoneurons were grown for 5-6 DIV on glass coverslips (0111500, Marienfeld GmbH) coated with PORN and laminin-111. For hippocampal neurons, cells were grown for 6 DIV on glass coverslips coated with poly-l-lysine and laminin-111. Cells were washed three times with RNase-free DPBS and fixed for 10 min in paraformaldehyde lysine phosphate (PLP) buffer (pH7.4) containing 4% paraformaldehyde (PFA) (28908, Thermo Fisher Scientific), 5.4% glucose and 0.01 M sodium metaperiodate. After three PBS washes, cells were permeabilized with the detergent solution provided with the kit for 4 min. Afterwards, the *Mapt* probe (VB1-13897-VC, Thermo Fisher Scientific) was diluted at 1:100 in the supplied buffer, and cells were incubated with the diluted probe overnight at 40°C. The following day, pre-amplifier, amplifier, and label probe solutions diluted 1:25 in the respective buffers were added to the cells sequentially for 1 h each at 40°C. Three washing steps were performed with the supplied wash buffer before each incubation and after the last one. Next, cells were washed twice with RNase-free DPBS followed by 4′,6-diamidino-2-phenylindole (DAPI)-staining and immunofluorescence labelling of Tubulin.

### Puromycin labelling-proximity ligation assay (Puro-PLA)

PLA was carried out using the Duolink InSitu Orange Starter Kit Mouse/Rabbit (DUO92102, Sigma-Aldrich) according to the manufacturer’s recommendations. Briefly, motoneurons grown for 6 DIV on laminin-111-coated glass coverslips were treated with 10 µg/mL puromycin (P8833, Sigma-Aldrich) for 10 min at 37°C. Cells were washed twice with DPBS, fixed in PLP for 10 min followed by permeabilization and washing. Cells were blocked in blocking buffer for 1 h at 37°C and incubated overnight at 4°C with anti-puromycin and anti-tau antibodies (Supplementary Table 1) diluted in blocking buffer. PLA probes were applied at 1:5 dilution for 1 h at 37°C followed by ligation for 30 min at 37°C and amplification for 100 min at 37°C. Cells were fixed again for 10 min at room temperature in PLP, washed with DPBS, and processed further for immunofluorescence and DAPI staining.

### Immunofluorescence staining

Motoneurons were cultured on laminin-111- and PORN-coated glass coverslips for 11 DIV. For hippocampal neurons, cells were cultured on laminin-111- and poly-L-lysine-coated glass coverslips for 22-25 DIV. Cells were washed twice with DPBS and fixed with 4% PFA at room temperature for 15 min followed by permeabilization with 0.1% Triton X-100 at room temperature for 20 min. After three washes in DPBS, cells were blocked in a blocking buffer containing 4% donkey serum at room temperature for 1 h. Primary antibodies diluted in blocking solution were applied onto coverslips and incubated at 4°C overnight followed by incubation with secondary antibodies at room temperature for 1 h and counterstaining with DAPI. A list of all antibodies is provided as Supplementary Table 1. Coverslips were washed and mounted using FluorSave Reagent (345789, Merck).

### Immunohistochemistry

Mice were deeply anesthetized and transcardially perfused with 0.1 M phosphate buffer (PB) (pH 7.4) and 4% PFA in PB. Brains were dissected and post-fixated overnight in PFA at 4°C. Samples were washed, embedded in 6% agarose cubes, and cut into 50 µm coronal sections using a vibratome (VT1000S, Leica). For antibody labelling, sections were washed in PBS, blocked with 4% donkey serum and 0.3% Triton X-100 in PBS for 2 h at room temperature, and subsequently incubated with the primary antibodies at 4°C for 48 h in blocking solution. After washing three times with PBS containing 0.3% Triton X-100, sections were incubated with secondary antibodies for 2 h at room temperature and washed again three times with PBS containing 0.3% Triton X-100. A list of all antibodies is provided as Supplementary Table 1. Then, sections were DAPI-stained, mounted onto object slides and embedded with FluorSave Reagent.

### Image acquisition and data analysis

Images were acquired on an Olympus Fluoview 1000 confocal system equipped with the following objectives: 10× (NA: 0.25), 20× (NA: 0.75), 40× (oil differential interference contrast, NA: 1.30), or 60× (oil differential interference contrast, NA: 1.35). Images were obtained with the Olympus FV10-ASW imaging software for visualization using 405, 473, 559, and 633 nm lasers. The resulting images (Olympus .oib format) were processed using ImageJ software as part of the Fiji package^39^. For FISH, maximum intensity projections were created from 0.3 μm z-stacks. For intensity measurements, raw images were projected using ImageJ and mean grey values were measured after background subtraction. For motoneurons, axonal tau intensity was measured in 20 µm-long proximal and distal regions and, for hippocampal neurons, in 20 µm-long distal regions. For axon length measurements, neurons were immunostained with anti-Tubulin antibodies and imaged on a Keyence BZ-8000K fluorescence microscope equipped with a standard colour camera using a 20× 0.7-NA objective. The length of the longest axon branch was quantified using ImageJ. Axon collaterals were not considered for the analysis. Brain sections were imaged on a fluorescence microscope (AxioImager 2, Zeiss) and z-stack images were adjusted for brightness and contrast using ImageJ software. 6E10, anti-Iba1 and AT8 immunosignals were quantified semiautomatically using the ImageJ threshold and particle analysis plugin after background subtraction.

### Quantitative PCR

qPCR reactions were set up with the Luminaris HiGreen qPCR Master Mix (K0994, Thermo Fisher Scientific) and run on a LightCycler 96 (Roche). qPCR primers were designed using the online Primer3Plus software. Primer sequences are listed in Supplementary Table 2. An annealing temperature of 60°C was used for all primers. Two technical qPCR replicates were set up for each sample and their Ct values were averaged before normalization and statistical analyses. Relative expression was calculated using the ΔΔCt method.

### Statistics and reproducibility

All statistical analyses were performed using GraphPad Prism version 9 for Windows (GraphPad Software, San Diego, California USA) and statistical significance was considered at test level *P*LJ<LJ0.05. Quantitative data are presented as meanLJ±LJs.d. unless otherwise indicated. Data for FISH, immunointensity and axon length measurements are represented as SuperPlots^40^. No statistical method was used to predetermine sample size. No data were excluded from the analyses. Two groups were compared using unpaired two-tailed Student’s *t*-tests or two-tailed one-sample *t*-tests. For multiple independent groups, one-way analysis of variance (ANOVA) with post hoc multiple comparisons test was used. Most experiments were carried out independently at least three times. Details of replicate numbers, quantification and statistics for each experiment are specified in the figure legends.

## Data availability

Sequences of MAPT-ASOs are provided in the Article.

## Acknowledgements

This work was supported by the Deutsche Forschungsgemeinschaft (BR4910/2–2 to M.B. and SE697/5–2 to M.S.) and a Grant from the Hermann und Lilli Schilling Stiftung im Stifterverband für die Deutsche Wissenschaft to M.S.

## Contributions

H.Z., M.B. and M.S. conceived the study and designed the experiments. M.B. designed the ASOs. H.Z. and S.S. performed all experiments. H.Z., S.S. and M.B. performed data analysis and interpretation. H.Z. and S.S. prepared figures. M.B. and M.S. supervised the study. M.B. wrote the manuscript with assistance from all authors.

## Additional information

### Supplementary information

Supplementary Tables 1-2.

### Corresponding authors

Correspondence to Michael Briese or Michael Sendtner.

### Extended data figure legends

**Extended Data Fig. 1: Comparison of MAPT-ASO1 and -ASO2 treatment for reduction of axonal *Mapt* mRNA levels.**

**a**, Immunofluorescence imaging of DIV 6 mouse motoneurons (MN) treated with different concentrations of a Cy3-labeled sense oligonucleotide. Scale bars: 10 µm and 5 µm (inset). **b**, Immunofluorescence imaging of DIV 25 untreated (Ctrl) mouse hippocampal neurons (HN) and hippocampal neurons treated with 10 µM of a Cy3-labeled sense oligonucleotide. Scale bars: 10 µm and 5 µm (inset). Images in **a** and **b** are representative of at least three independent experiments. **c**, *Mapt* FISH of DIV 6 untreated (Ctrl) mouse motoneurons and motoneurons treated with MAPT-ASO1 or -ASO2. Scale bars: 10 µm and 5 µm (inset). **d**, SuperPlots of the number of *Mapt* FISH punctae in the somata and axons of motoneurons. Statistical analysis was performed using a one-way ANOVA with Tukey’s multiple comparisons test. Data are mean ± s.d. of *n*LJ=LJ3 independent experiments. **e**, *Mapt* FISH of DIV 6 untreated (Ctrl) mouse hippocampal neurons and hippocampal neurons treated with MAPT-ASO1 or -ASO2. Scale bars: 10 µm and 5 µm (inset). **f**, SuperPlots of the number of *Mapt* FISH punctae in the somata and axons of hippocampal neurons. Statistical analysis was performed using a one-way ANOVA with Tukey’s multiple comparisons test. Data are mean ± s.d. of *n*LJ=LJ3 independent experiments.

**Extended Data Fig. 2: Comparison of MAPT-ASO1 and -ASO2 treatment for reduction of axonal tau levels.**

**a**, Tau immunostaining of DIV 11 untreated (Ctrl) mouse motoneurons and motoneurons treated with MAPT-ASO1 or -ASO2, with proximal and distal regions of the axon marked. Scale bars: 10 µm and 5 µm (inset). **b**, SuperPlots of tau immunointensities in the somata and proximal and distal axonal regions of untreated and treated motoneurons. Statistical analysis was performed using a one-way ANOVA with Tukey’s multiple comparisons test. Data are mean ± s.d. of *n*LJ=LJ3 independent experiments. **c**, Tau immunostaining of DIV 22 untreated (Ctrl) mouse hippocampal neurons and hippocampal neurons treated with MAPT-ASO1 or -ASO2. Scale bars: 10 µm and 5 µm (inset). **d**, SuperPlots of tau immunointensities in the somata and axons of untreated and treated hippocampal neurons. Statistical analysis was performed using a one-way ANOVA with Tukey’s multiple comparisons test. Data are mean ± s.d. of *n*LJ=LJ3 independent experiments.

**Extended Data Fig. 3: Treatment with MAPT-ASO2 reduces axonal tau levels in motoneurons.**

**a**, Immunofluorescence imaging of DIV 6 untreated (Ctrl) mouse motoneurons and motoneurons treated with 10 µM of a Cy3-labeled scramble oligonucleotide as control or MAPT-ASO2. Scale bars: 10 µm and 5 µm (inset). Images are representative of at least three independent experiments. **b**, *Mapt* FISH of DIV 6 untreated (Ctrl) mouse motoneurons and motoneurons treated with scramble oligonucleotide or MAPT-ASO2. Scale bars: 10 µm and 5 µm (inset). **c**, SuperPlots of the number of *Mapt* FISH punctae in the somata and axons of motoneurons. Statistical analysis was performed using a one-way ANOVA with Tukey’s multiple comparisons test. Data are mean ± s.d. of *n*LJ=LJ3 independent experiments. **d**, Tau immunostaining of DIV 11 untreated (Ctrl) mouse motoneurons and motoneurons treated with scramble oligonucleotide or MAPT-ASO2, with proximal and distal regions of the axon marked. Scale bars: 10 µm and 5 µm (inset). **e**, SuperPlots of tau immunointensities in the somata and proximal and distal axonal regions of motoneurons. Statistical analysis was performed using a one-way ANOVA with Tukey’s multiple comparisons test. Data are mean ± s.d. of *n*LJ=LJ5 independent experiments.

**Extended Data Fig. 4: Reduced axonal tau synthesis by MAPT-ASO2 treatment.**

**a**, Puromycin labelling with proximity ligation assay (Puro-PLA) for visualization of newly synthesized tau in DIV 6 untreated (Ctrl) mouse motoneurons and motoneurons treated with scramble oligonucleotide or MAPT-ASO2 using. Scale bars: 10 µm and 5 µm (inset). **b**, SuperPlots of tau Puro-PLA intensities in the somata and axons of motoneurons. Data are mean ± s.d. of *n*LJ=LJ2 independent experiments.

**Extended Data Fig. 5: Screening of additional ASOs for reducing axonal Mapt mRNA levels.**

**a**, Binding sites of MAPT-ASO1 to -ASO20 in the *Mapt* 3’ UTR. **b**, Sequences of MAPT-ASO1 to -ASO20 and their binding positions along human *MAPT* NCBI Reference Sequence NG_007398.2. *Mapt* mRNA levels were quantified by FISH in somata and axons of ASO-treated mouse hippocampal neurons and normalized to untreated hippocampal neurons. MAPT-ASOs with >50% reduction of axonal *Mapt* mRNA levels are highlighted in yellow. **c**, Quantification of the number of *Mapt* FISH punctae in the somata and axons of hippocampal neurons at DIV6. Data are shown as Tukey box plots.

