## Supplementary figures and images for "Prevention of tau accumulation through inhibition of hnRNP R-dependent axonal *Mapt* mRNA localization"

### Extended Data Fig. 1

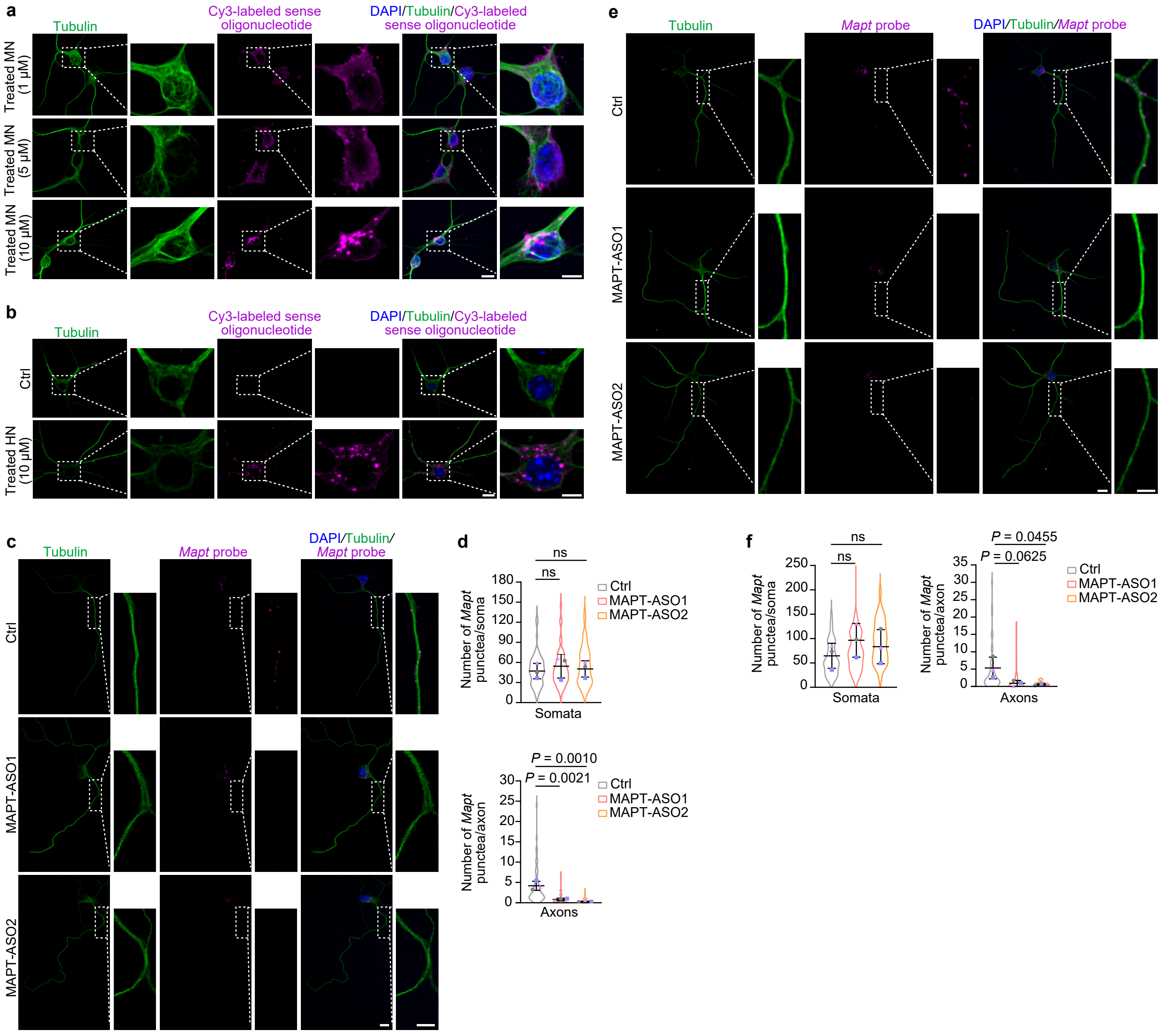

### Extended Data Fig. 2

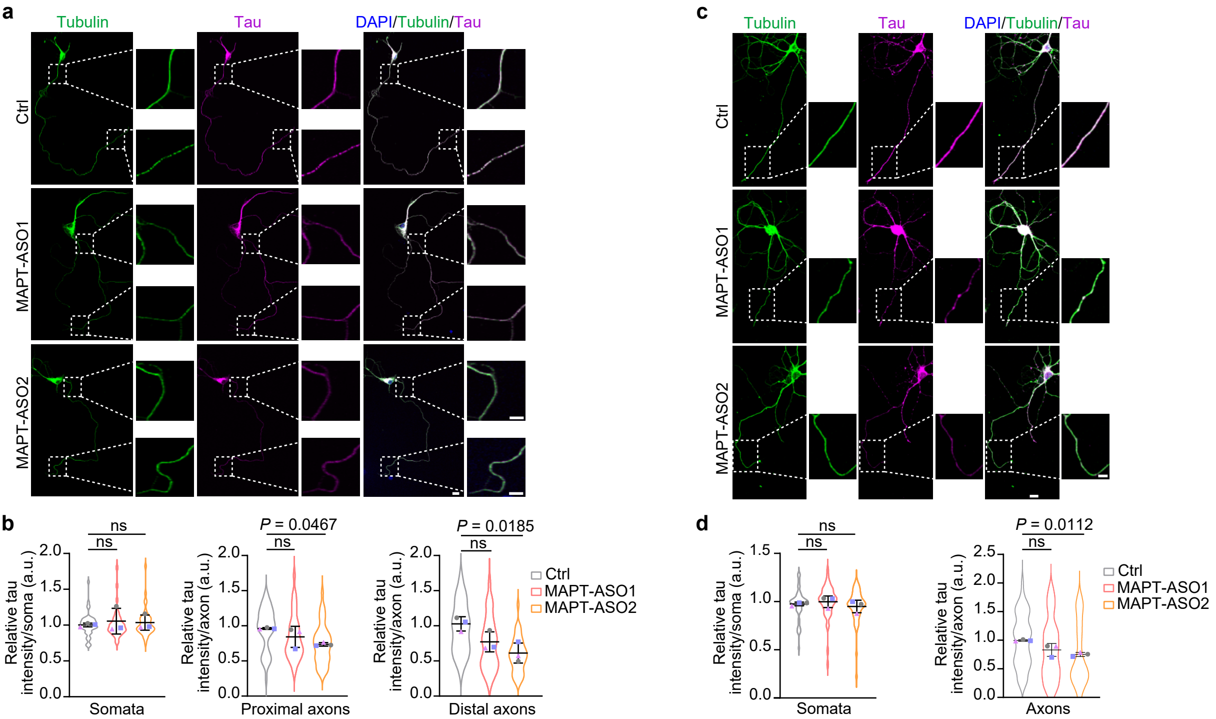

### Extended Data Fig. 3

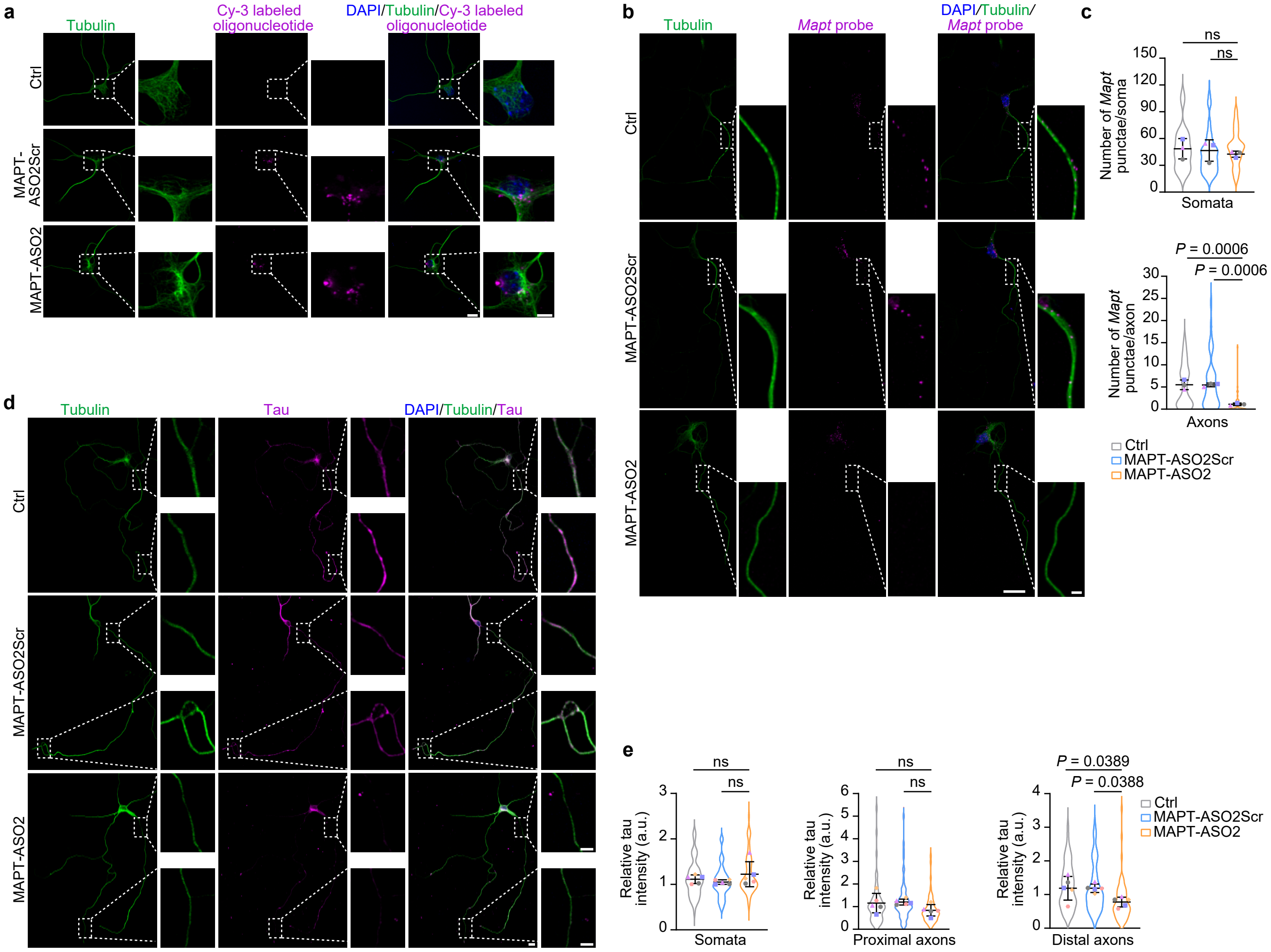

### Extended Data Fig. 4

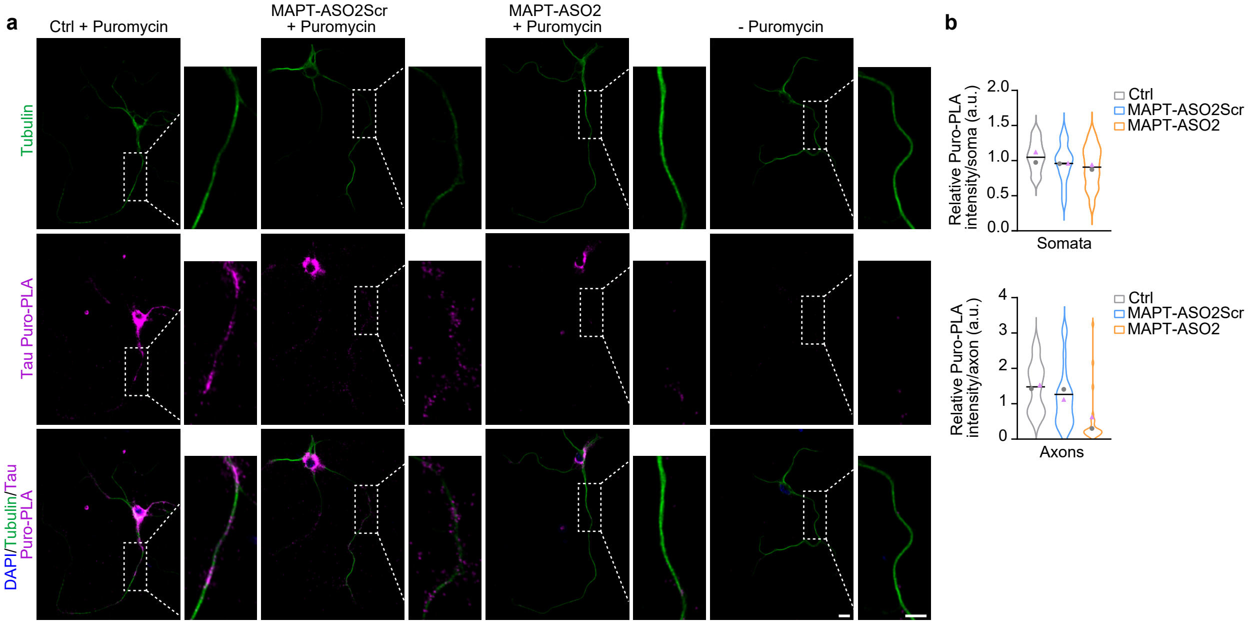
