## Extended Data Fig. 5 for "Prevention of tau accumulation through inhibition of hnRNP R-dependent axonal *Mapt* mRNA localization"

a

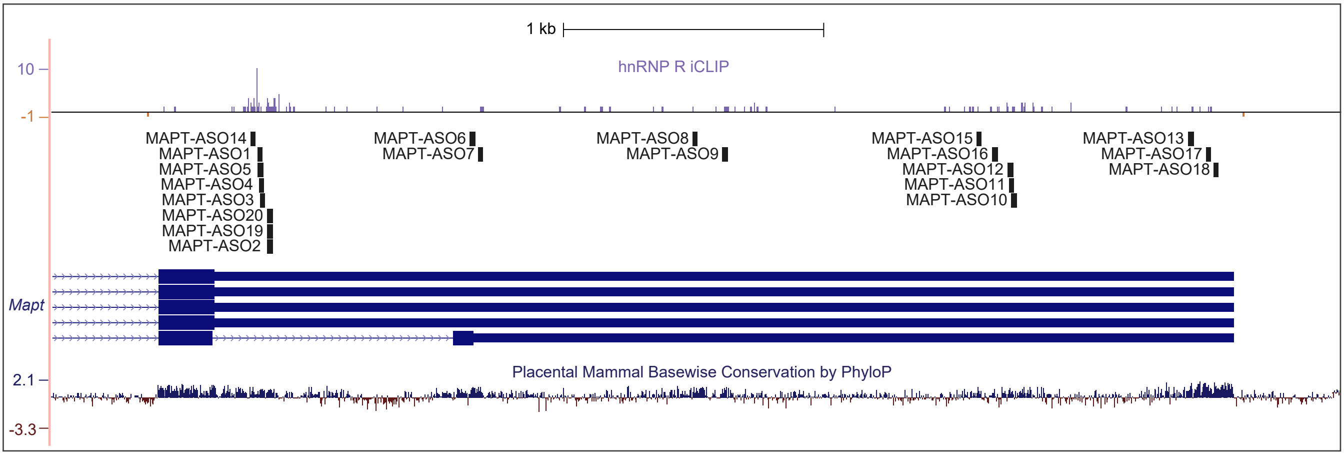


b

| ASO name | Sequence | Start position in NG_007398.2 | End position in NG_007398.2 | Relative *Mapt* expression soma | Relative *Mapt* expression axon |
|---|---|---|---|---|---|
| MAPT-ASO1 | TTTTGAAGTCCCGAGCCAAA | 134864 | 134883 | 1.69 | **0.40** |
| MAPT-ASO2 | TCAATTTGGAAAGATGAAAT | 134906 | 134925 | 1.38 | **0.31** |
| MAPT-ASO3 | TCCCATCACTGATTTTGAAG | 134876 | 134895 | 1.20 | **1.11** |
| MAPT-ASO4 | ATCACTGATTTTGAAGTCCC | 134872 | 134891 | 1.78 | **0.70** |
| MAPT-ASO5 | TGATTTTGAAGTCCCGAGCC | 134867 | 134886 | 1.26 | **0.17** |
| MAPT-ASO6 | TTAGCCCTAAAGTCCCAGGT | 135691 | 135710 | 1.17 | **0.66** |
| MAPT-ASO7 | CCAAGAGGCACAAGTCCTTA | 135724 | 135743 | 0.83 | **0.75** |
| MAPT-ASO8 | CAGACAAATCCAACTACAAC | 136566 | 136585 | 1.06 | **0.50** |
| MAPT-ASO9 | ATTTCAAGATACATGCGTCC | 136645 | 136664 | 1.14 | **0.94** |
| MAPT-ASO10 | TGAAGTCAATTTAAATGGAA | 138016 | 138035 | 1.23 | **0.81** |
| MAPT-ASO11 | TTTAAATGGAACTATTGATA | 138007 | 138026 | 1.49 | **0.32** |
| MAPT-ASO12 | TGGAACTATTGATAAAGTGA | 138001 | 138020 | 1.65 | **0.51** |
| MAPT-ASO13 | AAGAAATCATGGGACTTGCA | 138696 | 138715 | 1.31 | **0.72** |
| MAPT-ASO14 | ACAAAAGCAGGTTA | 134842 | 134855 | 0.77 | **0.41** |
| MAPT-ASO15 | GGGGGATTGTCCTCA | 137878 | 137892 | 0.65 | **0.38** |
| MAPT-ASO16 | CTATCTAGCCCACCCAA | 137949 | 137965 | 1.17 | **0.99** |
| MAPT-ASO17 | GACATTCACAGACAG | 138768 | 138782 | 1.06 | **0.47** |
| MAPT-ASO18 | ATCATTTGTTAAAACACA | 138797 | 138814 | 1.12 | **0.45** |
| MAPT-ASO19 | CAATTTGGAAAGATGAAA | 134907 | 134924 | 0.98 | **0.32** |
| MAPT-ASO20 | AATTTGGAAAGATGAA | 134908 | 134923 | 0.99 | **0.23** |

c

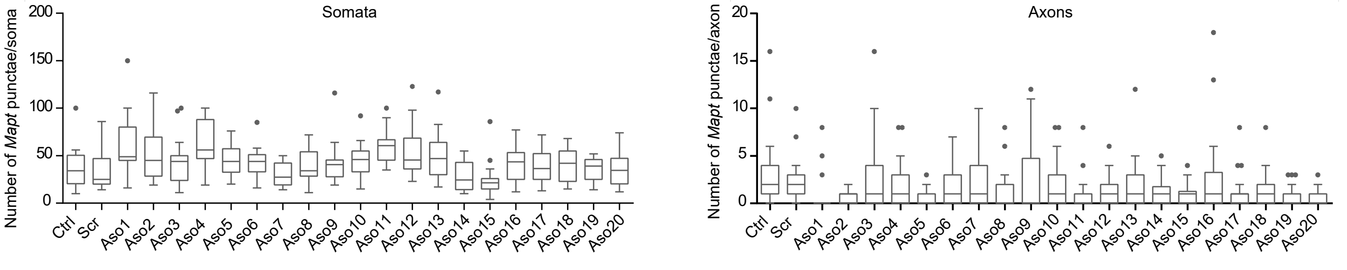
